# T cell responses to nonstructural protein 3 distinguish infections by Dengue and Zika viruses

**DOI:** 10.1101/305839

**Authors:** Bobby Brooke Herrera, Wen-Yang Tsai, Carlos Brites, Estela Luz, Celia Pedroso, Jan Felix Drexler, Wei-Kung Wang, Phyllis J. Kanki

**Affiliations:** Department of Immunology and Infectious Diseases, Harvard T.H. Chan School of Public Health, Boston, MA, USA; Department of Tropical Medicine, Medical Microbiology and Pharmacology, John A. Burns School of Medicine, University of Hawaii at Manoa, Honolulu, HI, USA; Laboratory of Infection Research, School of Medicine, Federal University of Bahia, Salvador, Brazil; Charité-Universitatsmedizin Berlin, corporate member of Freie Universität Berlin, Humboldt-Universität zu Berlin, and Berlin Institute of Health, Institute of Virology, Germany; German Centre for Infection Research, Germany

**Keywords:** Zika virus, dengue virus, T cell responses, nonstructural protein 3, Brazil

## Abstract

The 2015-16 Zika virus (ZIKV) epidemic in the Americas and the Caribbean demonstrates that clinical assays to detect, distinguish, and characterize immune responses to flaviviral infections are needed. ZIKV and dengue virus (DENV) are mosquito-transmitted flaviviruses sharing overlapping geographical distribution and have significant sequence similarity that can increase the potential for antibody and T cell cross-reaction. Using nonstructural protein 1-based enzyme-linked immunosorbent assays (ELISAs), we determine the serostatus of individuals living in a DENV- and ZIKV-endemic region in Brazil, identifying individuals with primary DENV (pDENV) and ZIKV (pZIKV), ZIKV with primary DENV (ZIKVwpDENV), and secondary DENV (sDENV) infections; pDENV and pZIKV were further confirmed by neutralization tests. Development of an enzyme-linked immunospot (ELISPOT) assay for DENV and ZIKV structural and nonstructural (NS) protein antigens enables us to distinguish infections by these viruses based on T cells and to characterize those responses. We find that IFN-γ and TNF-α T cell responses to NS3 differentiates DENV and ZIKV infections with 94% sensitivity and 92% specificity. In general, we also show that pDENV and sDENV cases and pZIKV and ZIKVwpDENV cases elicit similar T cell response patterns, and that HIV-infected individuals have T cell responses that are lower in magnitude compared to HIV-negative individuals. These results have important implications for DENV and ZIKV diagnostic and vaccine development and provide critical insights into the T cell response in individuals with multiple flaviviral infections.

## IMPORTANCE

The potential for antibody and T cell cross-reaction to DENV and ZIKV, flaviviruses that co-circulate and can sequentially infect individuals, has complicated diagnostic and vaccine development. Our serological data show that antibodies to nonstructural protein 1 can distinguish sequential human infections by DENV and ZIKV. The development of a simple and inexpensive assay also enables the differentiation of DENV and ZIKV infections based on the characterization of T cell responses. Our T cell data reveals strong response patterns that are similar in nature in individuals with one or multiple DENV infections and in individuals with only primary ZIKV infection and ZIKV-infected individuals with previous DENV exposure. The characterization of T cell responses in a serologically-validated group of individuals is of relevance to the development of vaccines and immunotherapeutics against these global threats.

## INTRODUCTION

*Aedes* mosquitoes transmit globally relevant flaviviruses including dengue virus (DENV) and Zika virus (ZIKV). DENV exists as four antigenic serotypes, DENV1 to DENV4 (1). These viruses have a wide geographic distribution with approximately 390 million infections annually and more than a quarter of the world’s population at risk (2). Prior to 2015, ZIKV was considered obscure and known to circulate in Africa and Southeast Asia as two separate viral lineages, African and Asian (3). While most asymptomatic, the clinical presentation of ZIKV infection resembles that of dengue including fever, rash, conjunctivitis, arthralgia, and myalgia (4). In early 2015, thousands of Asian ZIKV cases appeared in northeast Brazil, with accompanying reports of severe neuropathology including congenital microcephaly and Guillain-Barré syndrome (5, 6). In February 2016, the World Health Organization declared ZIKV a public health emergency of international concern (7). By June 2016, autochthonous transmission of ZIKV had been reported in 40 countries and territories throughout South and Central America and the Caribbean (8).

The emergence of ZIKV in DENV-endemic regions is of particular concern and relevant for diagnostic and vaccine development. The cocirculation of these genetically similar viruses can result in co-infection or sequential exposure, which has been shown to potentiate cross-reactive immunity at both the antibody and T cell levels (9–12). The envelope (E) protein is the major target of the antibody response in humans during flaviviral infection (1). Antibody-based assays were found to detect extensive crossreactivity to ZIKV E protein with other flaviviruses, requiring confirmation by plaque reduction neutralization tests (PRNTs) (11, 13–16). These tests, however, are challenged in their ability to confirm infection in individuals with multiple flaviviral infections especially during the acute and early convalescent phases. Several studies have also shown that most dengue-immune sera or DENV E monoclonal antibodies cross-react to ZIKV, but contain limited cross-neutralization activity and can instead enhance ZIKV infection, known as antibody-dependent enhancement (ADE) (17–22). In contrast, recent studies reported antibodies to ZIKV nonstructural protein 1 (NS1) were able to discriminate infections by these viruses (23, 24). We previously showed that combinations of DENV and ZIKV NS1-based enzyme-linked immunosorbent assays (ELISAs) were capable of distinguishing confirmed cases with past and present flaviviral infections including primary DENV (pDENV) and ZIKV (pZIKV), ZIKV with primary DENV (ZIKVwpDENV), and secondary DENV (sDENV) infections (12). These ELISAs are applicable for routine serological tests for DENV and ZIKV as well useful in retrospective studies to identify individuals with primary and multiple flaviviral infections.

Pre-existing T cell responses to DENV have also been shown to react to peptides encoded throughout the ZIKV proteome. DENV-naïve mice challenged with ZIKV developed ZIKV-specific CD8+ T cells, whereas DENV-immune mice challenged with ZIKV elicited cross-reactive CD8^+^ T cells that reduced infectious ZIKV (25). A study in humans infected with Asian ZIKV demonstrated that DENV serostatus influences the T cell response to ZIKV (10). DENV-immune individuals elicited CD4^+^ and CD8^+^ T cell responses to ZIKV more rapidly and of greater magnitude compared to DENV-naïve ZIKV-infected individuals. In addition, different patterns of immunodominant T cell responses were observed in the case of DENV and ZIKV infections. While CD8^+^ T cell responses against DENV target nonstructural (NS) proteins such as NS3, NS4B, and NS5, ZIKV-specific CD8^+^ T cell responses target the structural proteins, capsid (C), premembrane (prM), and E (10, 26). We previously developed a modified anthrax toxin (LFn)-based enzyme-linked immunospot (ELISPOT) assay, which revealed long-term T cell responses that were ZIKV- and DENV-specific to NS3 protease but cross-reactive to NS3 helicase in individuals infected with DENV and African ZIKV (27). The impact of cross-reactive immune responses in protection or development of ZIKV-mediated neuropathology remains unclear.

In this study, we utilized our NS1-based ELISAs to determine the DENV and ZIKV serostatus of individuals from Salvador, Brazil, a DENV-hyperendemic region with one of the highest incidence rates of ZIKV during the 2015-16 epidemic (28). We then tested the ability of our LFn ELISPOT assay to distinguish infections by DENV and Asian ZIKV based on T cells and to characterize those responses.

## RESULTS

### NS1-based ELISAs and neutralization test determine DENV and ZIKV serostatus

During the ZIKV outbreak in Salvador, Brazil, acute-phase blood samples were collected from hundreds of suspected ZIKV-infected patients attending HIV outpatient clinics between November 2015 and May 2016. Serological testing for ZIKV-NS1 IgG and DENV-E IgG was performed, revealing a high incidence of ZIKV infection in presumed DENV-immune and -naïve individuals (28). Fifty of these patients were included in the present study. Their median age was 43 (range: 23-72), 49% female, and 76% were human immunodeficiency virus (HIV)-infected. All HIV-infected individuals were on antiretroviral therapy; more than 92% had undetectable viral loads and normal CD4 counts. Patient characteristics and acute serology data are summarized in Table S1 and Table 1, respectively.

**Table 1.** Results of Serological tests

| ID | Acute-phase sera |  |  | Late convalescent-phase serum |  |  |  |  |  | Interpretation |
| --- | --- | --- | --- | --- | --- | --- | --- | --- | --- | --- |
|  | ZIKV-NS1 IgG<sup>a</sup> | DENV-E IgG<sup>a</sup> | PRNT test<sup>b</sup> | ZIKV-NS1 IgG<sup>c</sup> | DENV-NS1 IgG<sup>c</sup> | rOD ratio<sup>c</sup> | ZIKV-E IgG<sup>d</sup> | DENV-E IgG<sup>d</sup> | NT test<sup>f</sup><br>D1/D2/D3<br>/D4/ZIKV |  |
| ZK0978 | - | - | - | - | - | NA | - | - | <10/<10/<10/<10/<10 | negative |
| ZK0982 | - | - | - | - | - | NA | - | - | <10/<10/<10/<10/<10 | negative |
| ZK0987 | - | - | - | - | - | NA | - | - | <10/<10/<10/<10/<10 | negative |
| ZK0999 | - | - | - | - | - | NA | - | - | <10/<10/<10/<10/<10 | negative |
| ZK0979 | + | + |  | + | - | NA | +<sup>e</sup> | + | <10/<10/<10/<10/>160 | pZIKV |
| ZK0993 | + | + |  | + | - | NA | +<sup>e</sup> | + | <40/<40/<10/<10/>160 | pZIKV |
| ZK0998 | + | + |  | + | - | NA | +<sup>e</sup> | + | <10/<10/<40/<10/>160 | pZIKV |
| ZK1006 | + | + |  | + | - | NA | +<sup>e</sup> | + | <80/<40/<10/<40/640 | pZIKV |
| ZK0996 | + | - | + | + | - | NA | +<sup>e</sup> | + | <10/<10/<10/<10/>160 | pZIKV |
| ZK0966 | - | + | - | - | - | NA | + | +<sup>e</sup> | <40/160/<10/<40/<10 | pDENV |
| ZK0980 | - | + |  | - | - | NA | + | +<sup>e</sup> | <40/<40/>160/<40/<10 | pDENV |
| ZK0995 | - | + |  | - | - | NA | + | +<sup>e</sup> | >640/160/<10/<10/<10 | pDENV |
| ZK0997 | ND | ND |  | - | - | NA | + | +<sup>e</sup> | <40/>160/<40/<40/<10 | pDENV |
| ZK0972 | + | + |  | + | + | ≥0.24 | + | + | >40/ND/ND/ND/>80 | ZIKVwpDENV |
| ZK0975 | + | + |  | + | + | ≥0.24 | + | + | >40/ND/ND/ND/>80 | ZIKVwpDENV |
| ZK0989 | + | + | + | + | + | ≥0.24 | + | + | >40/ND/ND/ND/>80 | ZIKVwpDENV |
| ZK0991 | + | + | + | + | + | ≥0.24 | + | + | >40/ND/ND/ND/>80 | ZIKVwpDENV |
| ZK1000 | + | + |  | + | + | ≥0.24 | + | + | >40/ND/ND/ND/>40 | ZIKVwpDENV |
| ZK1009 | + | + | + | + | + | ≥0.24 | + | + | >10/ND/ND/ND/>80 | ZIKVwpDENV |
| ZK1011 | + | + |  | + | + | ≥0.24 | + | + | >40/ND/ND/ND/>80 | ZIKVwpDENV |
| ZK1012 | + | + | + | + | + | ≥0.24 | + | + | >40/ND/ND/ND/>80 | ZIKVwpDENV |
| ZK1014 | + | + | + | + | + | ≥0.24 | + | + | >40/ND/ND/ND/>80 | ZIKVwpDENV |
| ZK1015 | + | + | + | + | + | ≥0.24 | + | + | >40/ND/ND/ND/>80 | ZIKVwpDENV |
| ZK0968 | + | ND |  | + | + | ≥0.24 | + | + | >40/ND/ND/ND/>80 | ZIKVwpDENV |
| ZK0984 | + | + |  | + | + | ≥0.24 | + | + | >320/>1280/<80/>1280/>320 | ZIKVwpDENV |
| ZK0976 | - | + |  | + | + | <0.24 | + | + | >40/ND/ND/ND/<10 | sDENV |
| ZK0986 | - | + |  | + | + | <0.24 | + | + | >40/ND/ND/ND/>80 | sDENV |
| ZK0967 | + | + |  | + | + | <0.24 | + | + | >40/ND/ND/ND/>80 | sDENV |
| ZK0969 | + | + |  | + | + | <0.24 | + | + | >40/ND/ND/ND/>40 | sDENV |
| ZK0971 | + | + |  | + | + | <0.24 | + | + | >40/>40/ND/ND/<10 | sDENV |
| ZK0977 | + | + |  | + | + | <0.24 | + | + | >40/ND/ND/ND/>80 | sDENV |
| ZK0983 | + | + | + | + | + | <0.24 | + | + | >40/ND/ND/ND/>10 | sDENV |
| ZL0985 | + | + | + | + | + | <0.24 | + | + | >40/ND/ND/ND/>40 | sDENV |
| ZK0988 | + | + |  | + | + | <0.24 | + | + | >40/ND/ND/ND/>40 | sDENV |
| ZK0992 | + | + | + | + | + | <0.24 | + | + | >40/ND/ND/ND/>80 | sDENV |
| ZK0994 | + | + | - | + | + | <0.24 | + | + | >40/ND/ND/ND/>40 | sDENV |
| ZK1001 | + | + |  | + | + | <0.24 | + | + | >40/>40/ND/ND/<10 | sDENV |
| ZK1010 | + | + |  | + | + | <0.24 | + | + | >40/ND/ND/ND/>80 | sDENV |
| ZK1013 | + | ND |  | + | + | <0.24 | + | + | >40/ND/ND/ND/>10 | sDENV |
| ZK0973 | - | ND |  | - | + | <0.24 | + | + | >40/<80/>80/ND/>320 | sDENV |
| ZK0974 | - | + |  | - | + | <0.24 | + | + | >40/>320/>80/ND/>10 | sDENV |
| ZK1002 | - | + | - | - | + | <0.24 | + | + | >40/>80/<80/ND/<10 | sDENV |
| ZK1003 | - | + | - | - | + | <0.24 | + | + | >40/>80/<80/ND/<10 | sDENV |
| ZK1007 | - | + |  | - | + | <0.24 | + | + | >40/>80/>80/ND/<10 | sDENV |
| ZK1016 | - | + |  | - | + | <0.24 | + | + | >40/>1280/>1280/ND/>10 | sDENV |
| ZK0990 | - | + | - | - | + | <0.24 | + | + | >40/>1280/>80/ND/<10 | sDENV |
| ZK1004 | + | + |  | - | + | NA | + | + | ND/ND/ND/ND/ND | unknown |
| ZK1005 | + | ND |  | - | + | NA | + | + | ND/ND/ND/ND/ND | unknown |
| ZK1008 | + | + |  | - | - | NA | + | + | ND/ND/ND/ND/ND | unknown |
| ZK0981 | - | ND |  | - | - | NA | - | - | ND/ND/ND/ND/ND | unknown |

pZIKV: primary ZIKV infection; pDENV: primary DENV infection; sDENV: secondary DENV infection; ZIKV/wpDENV: ZIKV infection
with previous DENV infection. ND: not determined; NA not applicable.
<sup>a</sup>Euroimmun ZIKV-NS1 and DENV-E IgG ELISAs were performed on acute-phase sera (28).
<sup>b</sup>PRNT was performed on acute-phase sera to detect neutralization antibody to ZIKV (47).
<sup>c</sup>ZIKV-NS1 and DENV-NS IgG ELISAs were described previously (12). The rOD ratio (ZIKV-NS1/DENV-NS1) < or ≥0.24 was classified as sDENV or ZIKV/wpDENV (12).
<sup>d,e</sup>ZIKV-E and DENV-E IgG ELISAs utilized ZIKV VLP and DENV virions, respectively (46). ΔrOD (rOD of ZIKV–rOD of DENV) ≥ 0.17 or < –0.17 was classified as pZIKV or pDENV infection, respectively.
<sup>f</sup>Mico-neutralization test (NT) were performed (with NT<sub>90</sub> titers shown) to confirm none, pZIKV or pDENV infection (46, 48).

In order to determine the DENV and ZIKV serostatus among the study participants who had potentially been dual exposed, we collected late convalescent-phase blood samples and employed our previously developed ZIKV-NS1 and DENV-NS1 IgG ELISAs (22). For samples positive for DENV-NS1, we calculated the ratio of relative optical density (rOD) of ZIKV-NS1 to that of DENV-NS1 and used the rOD ratio < or ≥ 0.24 to determine sDENV or ZIKVwpDENV infection, respectively (22). Twelve ZIKVwpDENV and 21 sDENV infections were identified (Table 1, Fig. 1A to C). Five samples with ZIKV-NS1 positive and DENV-NS1 negative were pZIKV. Since these samples were collected more than one year post-infection, some anti-NS1 antibodies may have declined to levels below detection, we further tested with ZIKV and DENV E protein-based ELISAs and identified four samples negative for both ZIKV and DENV in all four ELISAs tested (Table 1). Based on the difference in rOD of ZIKV and DENV E proteins (ΔrOD=rOD of ZIKV – rOD of DENV), we identified four pZIKV (ΔrOD ≥0.17) and four pDENV (ΔrOD <−0.17) infections (Table 1, Fig. 1D to E). The negative, pZIKV and, pDENV samples were further confirmed by micro-neutralization tests to DENV1-4 and ZIKV; all four negative samples had NT_90_ titers <10 to DENV1-4 and ZIKV, and the five pZIKV and four pDENV samples showed monotypic neutralization pattern to ZIKV and to one of the four DENV serotypes, respectively (Table 1). For the remaining 33 samples, microneutralization tests to ZIKV, DENV1 and/or DENV2 or 3 were performed to show that all 12 ZIKVwpDENV samples neutralize (NT_90_ titers ≥10) ZIKV plus at least one DENV serotype, whereas all 21 sDENV samples neutralize at least two DENV serotypes or DENV plus ZIKV; these patterns were compatible with unspecified flavivirus infection according to CDC guidelines (16). An additional three samples (ZK1004, 1005, 1008), which had ΔrOD between −0.17 and 0.17, positive ZIKV-NS1 IgG at acute-phase sera but negative at late convalescent-phase, were classified as undetermined (Table 1). Another sample (ZK0981), for which DENV acute-phase serology was not performed, was negative for ELISAs using late convalescent-phase serum and was also classified as undetermined.

**Figure 1.**
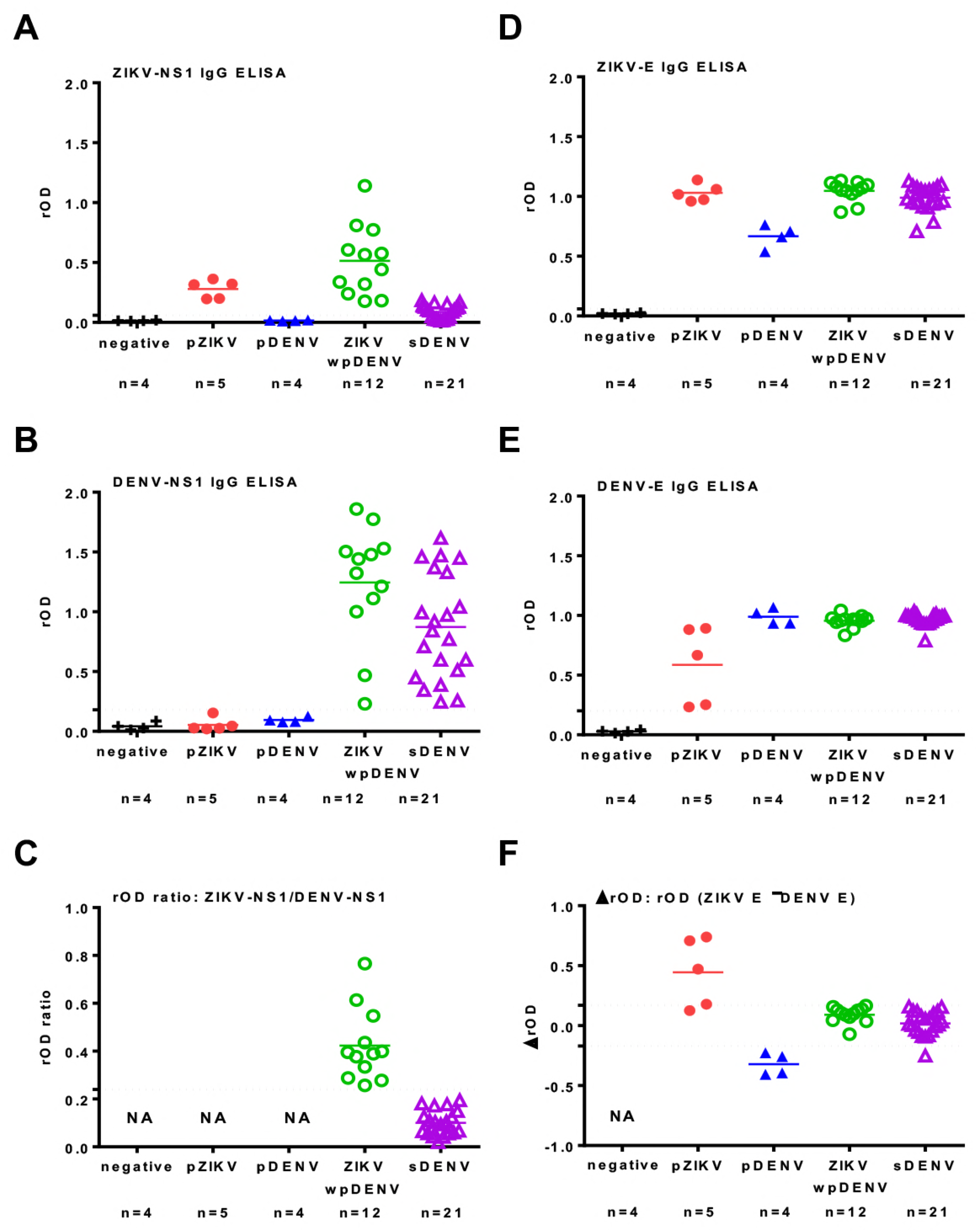
ZIKV and DENV NS1-based and E-based IgG ELISAs. (A) ZIKV-NS1, (B) DENV-NS1 IgG ELISAs and (C) rOD ratio. (D) ZIKV-E, (E) DENV-E and (F) ΔrOD=rOD of ZIKV-rOD of DENV. Dots lines indicate cut-off values, 0.24 for rOD ratio and 0.17 for ΔrOD. pZIKV: primary ZIKV infection; pDENV: primary DENV infection; sDENV: secondary DENV infection; ZIKVwpDENV: ZIKV infection with previous DENV infection. NA not applicable.

### T cell responses to NS3 distinguish DENV and ZIKV infections

We recently reported the development of a LFn ELISPOT assay based on NS3 protease and helicase to distinguish DENV and African ZIKV human infections (27). To assess the ability of the assay to distinguish infection by DENV and Asian ZIKV, we performed DENV and ZIKV homologous and heterologous LFn-NS3 protease and helicase stimulation of date-matched late convalescent-phase PBMCs in an IFN-γ and TNF-α ELISPOT among the serological-validated pDENV, pZIKV, sDENV, and ZIKVwpDENV, and undetermined cases. Using a NS3 protease to helicase ratio cutoff of 1.05 for the IFN-γ ELISPOT, pDENV and sDENV cases and pZIKV, ZIKVwpDENV and the 3 out of the 4 serologically undetermined cases appeared to group together (Fig. 2A). From the undetermined cases, 3 out of 4 (ZK1004, 1005, 1008) grouped with the ZIKV-exposed individuals, while ZK0981 grouped with the DENV-infected individuals. Using a ratio cutoff of 1.048 for the TNF-α ELISPOT, similar groupings were observed (Fig. 2B). We were unable to distinguish sequential infections based on T cell responses to NS3 protease and helicase.

**Figure 2.**
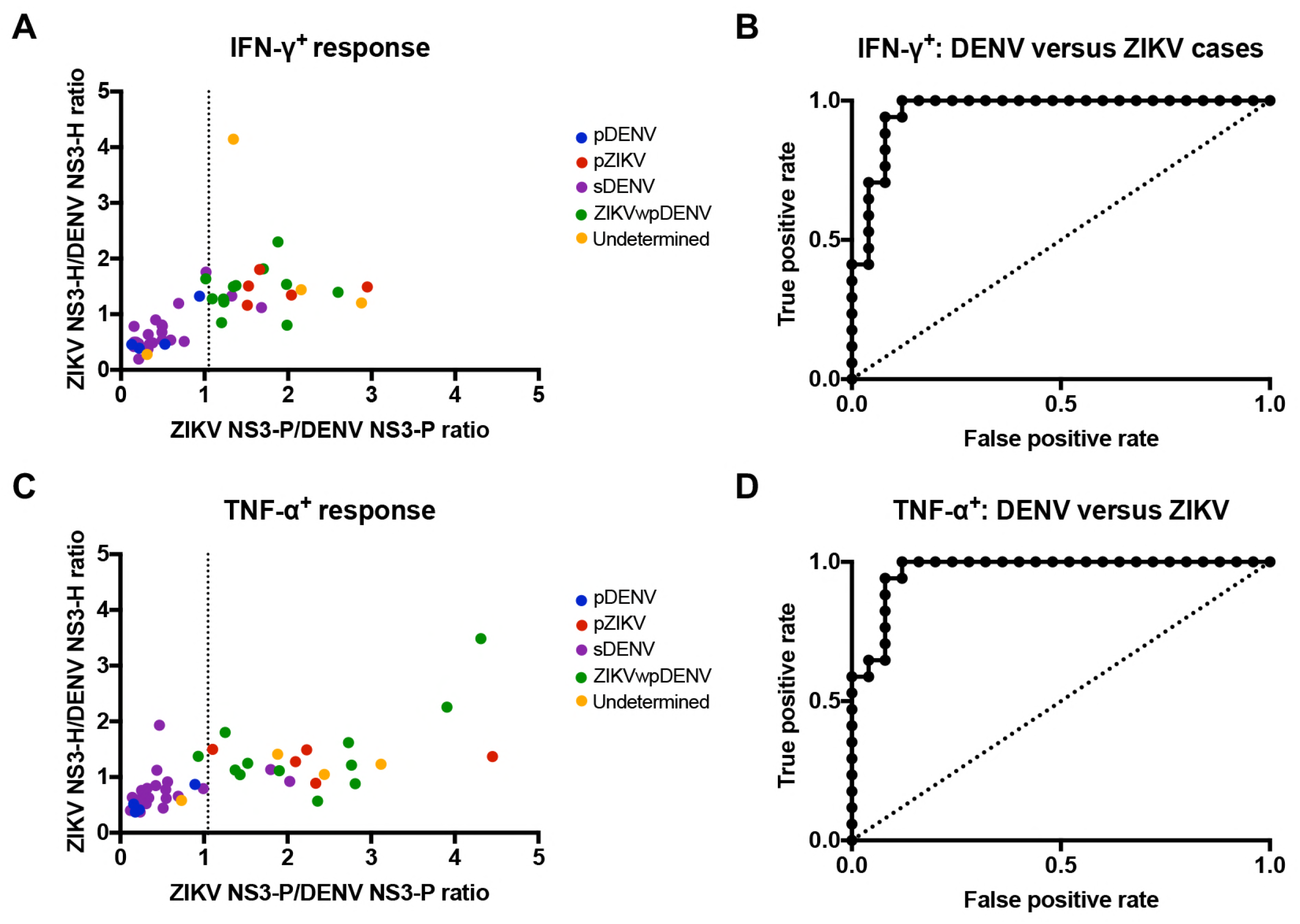
T cell responses to NS3 protease and helicase and ROC analysis of the ELISPOT test. Late convalescent-phase PBMCs from DENV- and/or ZIKV-infected individuals were treated with homologous and heterologous LFn-DENV and -ZIKV NS3 protease and the specific IFN-γ and TNF-α T cell responses were detected by *ex vivo* ELISPOTs. (A) Scatter plot of the ratios of ZIKV NS3 protease to DENV NS3 protease IFN-γ responses versus ratios of helicase. (B) ROC analysis for the IFN-γ ELISPOT. (C) Scatter plot of the ratios of ZIKV NS3 protease to DENV NS3 protease TNF-α T cell responses versus ratios of helicase. (D) ROC analysis for the TNF-α ELISPOT. The dashed line on (A) represents the optimal cutoff value of 1.05 and the dashed line on (C) represents the optimal cutoff value of 1.048. Individual colored dots represent serologically-validated DENV- and/or ZIKV-infected individuals and the undetermined cases.

Test data were further analyzed to define sensitivity (identifying true positives; individuals who had been infected by DENV versus ZIKV) and specificity (true negatives; DENV- or ZIKV-uninfected individuals). We evaluated sensitivity and specificity as functions of the IFN-γ and TNF-α cutoff values, above which a sample was considered positive and below which a sample was considered negative. We grouped pDENV and sDENV and pZIKV and ZIKVwpDENV cases together based on the clustering observed and excluded the 4 serologically undetermined cases from the analysis. Receiver Operating Characteristic (ROC) curves and corresponding numerical values illustrate the performance of the ELISPOTs as a function of the discrimination threshold, plotted as sensitivity versus 1 – specificity. The areas of the ROC curves represent test performance, where 1 represents a perfect test, and 0.5 represents a random predictor. We measured areas of 0.96 and 0.97 for the IFN-γ and TNF-α ELISPOTs, respectively (Table 2, Fig. 2 C-D). Using the cutoff values, the test sensitivity and specificity for both the IFN-γ and TNF-α ELISPOTs were 94% and 92%, respectively.

**Table 2.**
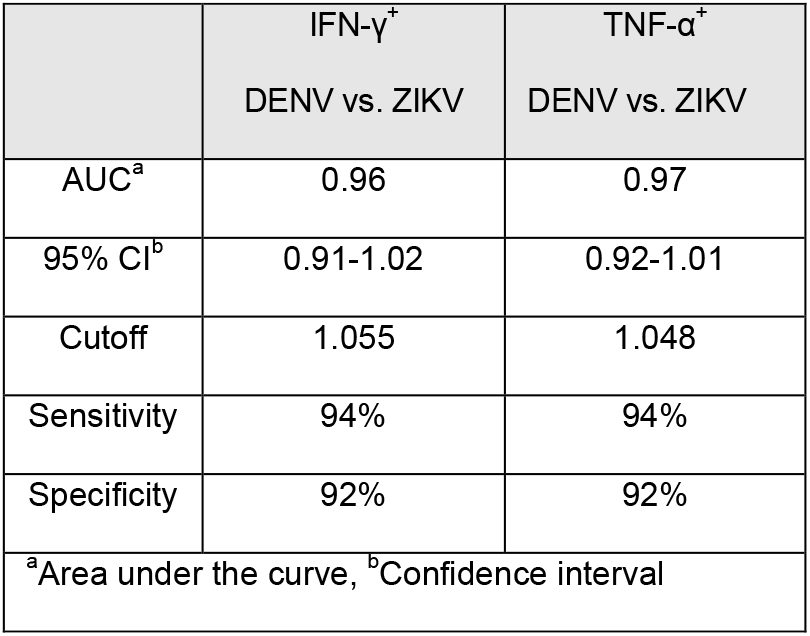
Numerical values of ROC analysis and sensitivity and specificity results

### LFn-DENV and -ZIKV structural and nonstructural proteins elicit robust T cell responses and prior DENV exposure does not affect the response

To assess the magnitude of T cell responses among the study participants, we stimulated late convalescent-phase PBMCs in IFN-γ and TNF-α ELISPOTs using the following six LFn fusion proteins: LFn-DENV-NS3-Protease (LFn-DV NS3-P), LFn-DENV-NS3-Helicase (LFn-DV NS3-H), LFn-ZIKV-Capsid (LFn-ZV C), LFn-ZIKV-premembrance (LFn-ZV prM), LFn-ZIKV-NS3-Protease (LFn-ZV NS3-P), LFn-ZIKV-NS3-Helicase (LFn-ZV NS3-H). Individuals with pDENV and sDENV infections elicited similar IFN-γ and TNF-α T cell response patterns (Fig. 3 A-B). These individuals had T cell responses to LFn-DV NS3-H and LFn-ZV NS3-H that were stronger in magnitude than to LFn-DV NS3-P and LFn-ZV NS3-P, respectively. Additionally, T cell responses to LFn-DV NS3-P and NS3-H were stronger compared to LFn-ZV NS3-P and NS3-H. The amount of T cell cross-reaction to the ZIKV structural proteins (LFn-ZV C and LFn-ZV prM) was limited, compared to high cross-reactivity to LFn-ZV NS3-P and NS3-H. Furthermore, individuals with pZIKV and ZIKVwpDENV infections elicited T cell responses to LFn-ZV NS3-H and LFn-DV NS3-H that were stronger in magnitude compared to LFn-ZV NS3-P and LFn-DV NS3-P, respectively (Fig. 3 C-D). While individuals with pZIKV and ZIKVwpDENV infections had stronger IFN-γ T cell responses to LFn-ZV NS3-H than to the ZIKV structural proteins, TNF-α responses to the ZIKV structural proteins were stronger than to LFn-ZV NS3-P.

**Figure 3.**
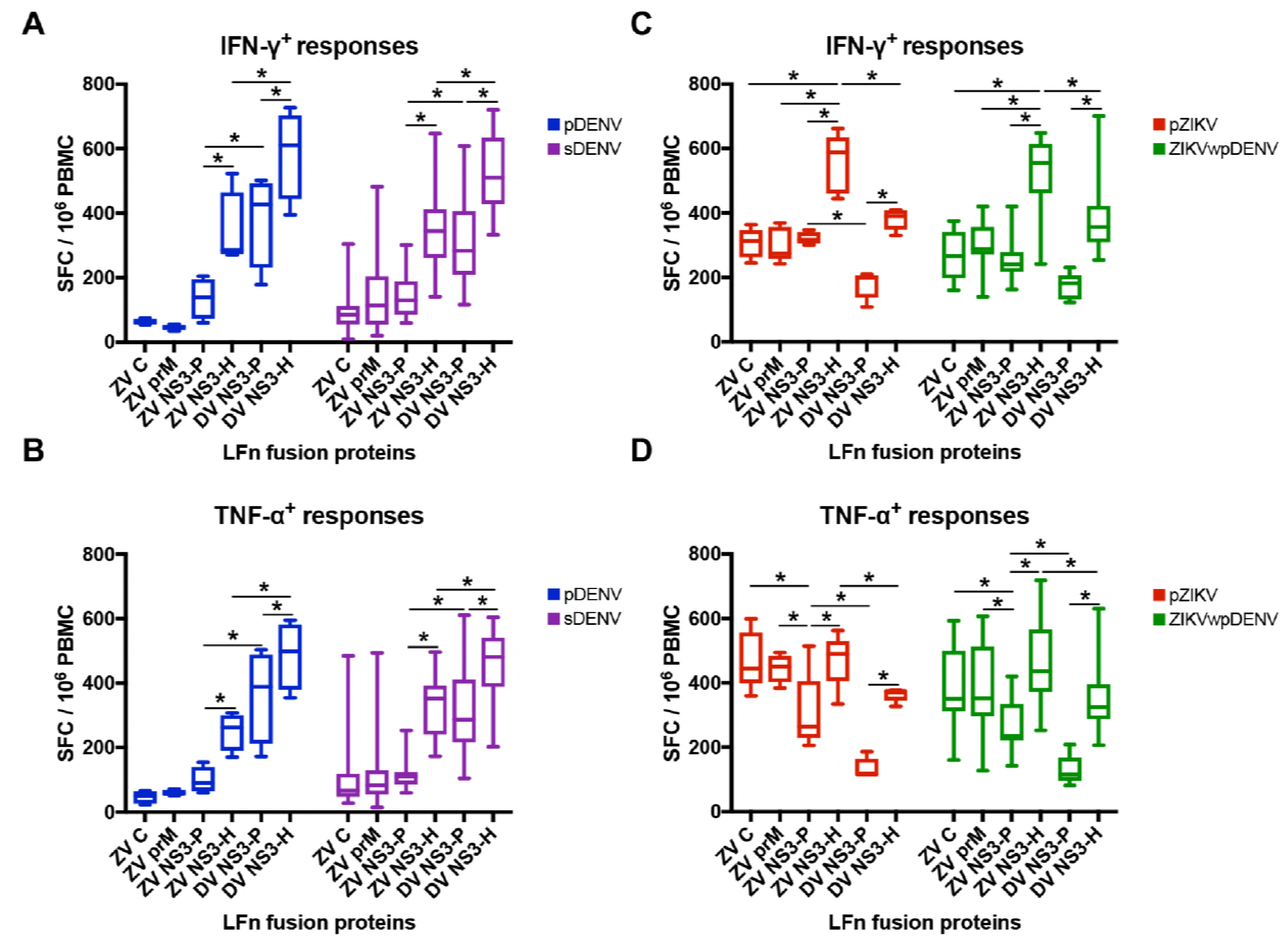
T cell responses to ZIKV and/or DENV structural or nonstructural proteins among subgroups with different DENV and ZIKV serostatus. Late convalescent-phase PBMCs from DENV- and/or ZIKV-infected individuals were treated with homologous and/or heterologous LFn-DENV and -ZIKV capsid (ZV C), premembrane (ZV prM), NS3 protease (DV or ZV NS3-P) and NS3 helicase (DV or ZV NS3-H) and the specific IFN-γ and TNF-α T cell responses were detected by *ex vivo* ELISPOTs. IFN-γ and TNF-α spot forming cells (SFC) were detected, counted, and expressed as box plots with mean and standard deviations. Comparison of late convalescent-phase (A) IFN-γ and (B) TNF-α T cell responses between individuals with pDENV and sDENV infections. Comparison of late convalescent-phase (C) IFN-γ and (D) TNF-α T cell responses between individuals with pZIKV and ZIKVwpDENV infections. Individual colored plots represent serologically-validated DENV- and/or ZIKV-infected individual. *, p<0.05.

We further evaluated the impact of DENV immunity on the magnitude of T cell responses. We compared the magnitude of the IFN-γ and TNF-α T cell responses between individuals with pDENV and sDENV infections and pZIKV and ZIKVwpDENV infections. In all cases, T cell responses in individuals with prior DENV exposure were not significantly higher compared to individuals with a primary DENV or ZIKV infection (Fig. 3 A-D). While IFN-γ and TNF-α responses appeared stronger to the ZIKV structural proteins in individuals with sDENV infections than to individuals with pDENV infections, these differences were not statistically significant. Similarly, individuals with ZIKVwpDENV infection had comparable IFN-γ and TNF-α T responses to those with pZIKV infections (Fig. 3 C-D).

### HIV influences the T cell response in DENV-exposed individuals

We also compared the magnitude of the IFN-γ and TNF-α T cell responses in DENV-exposed (grouping individuals with pDENV and sDENV infections together), pZIKV, and ZIKVwpDENV individuals who were HIV-negative or HIV-infected. DENV-exposed HIV-negative individuals had stronger IFN-γ responses to LFn-ZV C, -ZV NS3-P, and -ZV NS3-H compared to HIV-infected individuals. IFN-γ responses to LFn-DV NS3-P and NS3-H appeared to be stronger in the HIV-infected individuals, although these differences were not statistically significant (p=0.61 and p=0.13, respectively) (Fig. 4 A). A similar pattern of responses was observed for TNF-α (Fig. 4 D). In general, ZIKVwpDENV HIV-negative individuals had stronger IFN-γ and TNF-α responses compared to individuals that were HIV-infected (Fig. 4 B and E). In contrast, there was largely no difference in the IFN-γ and TNF-α responses in pZIKV HIV-negative and HIV-infected individuals (Fig. 4 C and F). There was an exception where the TNF-α response to LFn-ZV NS3-H was stronger in individuals that were HIV-negative.

**Figure 4.**
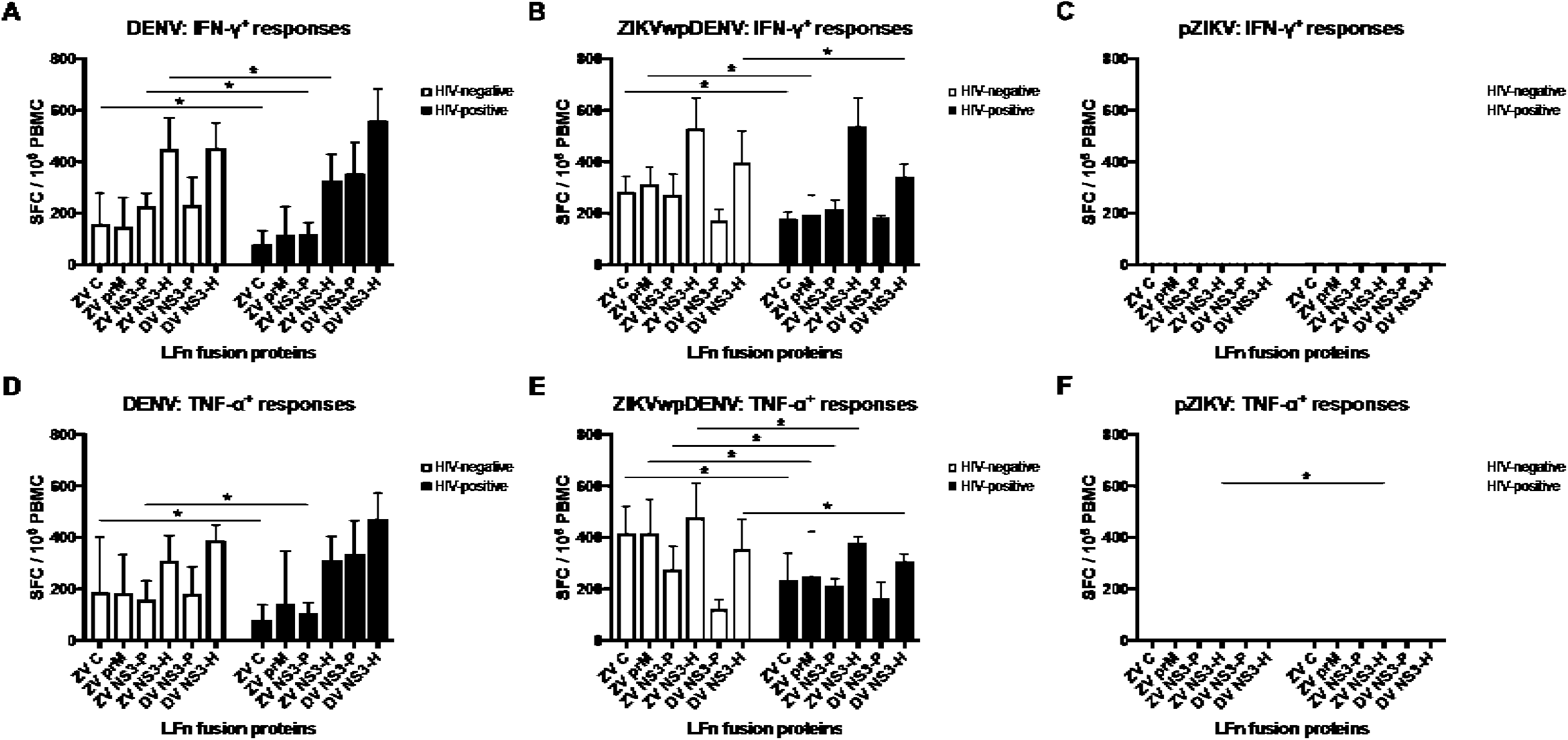
Impact of HIV status on the T cell response. Comparison of mean convalescent-phase IFN-γ T cell responses expressed as bars and standard deviation between (A) HIV-negative (open black bars) and HIV-infected (shaded black bars) individuals with pDENV and sDENV infections grouped together, (B) HIV-negative (open dark grey bars) and HIV-infected (shaded dark grey bars) individuals with ZIKVwpDENV infections, and (C) HIV-negative (open light grey bars) and HIV-infected (shaded light grey bars) individuals with pZIKV infections. Comparison of mean convalescent-phase TNF-α T cell responses expressed as bars and standard deviation between (D) HIV-negative (open black bars) and HIV-infected (shaded black bars) individuals with pDENV and sDENV infections grouped together, (E) HIV-negative (open dark grey bars) and HIV-infected (shaded dark grey bars) individuals with ZIKVwpDENV infections, and (F) HIV-negative (open light grey bars) and HIV-infected (shaded light grey bars) individuals with pZIKV infections. *, p<0.05.

## DISCUSSION

We report on the characterization of late convalescent-phase antibody and T cell responses in individuals from Salvador, Brazil, a DENV-hyperendemic region that was burdened by the 2015-2016 ZIKV outbreak. Our study presents three major findings in a serologically-validated group of DENV and/or ZIKV infected individuals. First, IFN-γ and TNF-α T cell response ratios of ZIKV NS3 protease to DENV NS3 protease can discriminate infections in individuals exposed to these viruses. Second, individuals with pDENV and sDENV infections have similar T cell response patterns, with extensive cross-reactivity to ZIKV NS3 helicase, whereas individuals with pZIKV and ZIKVwpDENV infections have strong responses to both ZIKV structural and nonstructural proteins, with high cross-reaction to DENV NS3 helicase. Third, HIV-infection is associated with responses that are lower in magnitude in DENV exposed individuals.

Our previous study of NS1-based ELISAs on convalescent-phase sera from RT-PCR confirmed cases with pZIKV, pDENV, sDENV and ZIKVwpDENV infections showed that sDENV infection panel cross-react to ZIKV-NS1 and the rOD ratio of ZIKV-NS1 to DENV-NS1 in IgG ELISA can distinguish sDENV and ZIKVwpDENV infections (22). Since anti-NS1 antibodies may decline over time and become undetectable especially for those with primary infection, we further tested these samples with E protein-based IgG ELISAs and identified four negative samples, five pZIKV and four pDENV infections. All these 13 samples have been verified by neutralization test using NT_90_ ≥10 as cutoff based on the CDC guidelines (16), suggesting that ΔrOD based on ZIKV-E and DENV-E IgG ELISAs can distinguish pZIKV and pDENV infections; this could potentially be a useful tool for epidemiology and pathogenesis study in endemic regions. However, the sample size is small and the ΔrOD of 0.17 was based on a single serum dilution of 1:800, future studies involving larger sample size and different dilutions or end-point titers are needed to further validate these observations.

The degree of amino acid sequence identity between DENV and ZIKV structural and nonstructural proteins is 49% and 51%, respectively (10). Multiple sequence alignment and homology determination of DENV and ZIKV NS3 demonstrates high amino acid sequence identity of 67%, with protease and helicase homology of 58% and 72%, respectively, consistent with the higher degree of DENV/ZIKV cross-reaction in NS3 helicase (Table 3, Fig. 5). Our recent characterization of acute- and convalescent-phase T cells collected from individuals infected with DENV and African ZIKV in Senegal, West Africa, revealed sustained DENV- and ZIKV-specific responses to NS3 protease and cross-reactive responses to NS3 helicase (27). Our findings in individuals infected with DENV and Asian ZIKV are in agreement with our previous observations. Although we were unable to distinguish sequential exposure, the LFn NS3 protease ELISPOT differentiates infections between DENV- and ZIKV-infected individuals with high sensitivity and specificity of 94% and 92%, respectively.

**Table 3.** Sequence homology of DENV and Asian ZIKV NS3^a^

|  | % homology to ZIKV |  |  |
| --- | --- | --- | --- |
| Serotype | NS3 protease | NS3 helicase | Full-length |
| DENV1 | 55% | 71% | 66% |
| DENV2 | 58% | 72% | 67% |
| DENV3 | 58% | 72% | 67% |
| DENV4 | 59% | 71% | 67% |
| <sup>a</sup>Homology analysis between Asian ZIKV (GenBank accession number: NC_035889.1) and DENV1 (ACO06157.1), DENV2 (JN819419.1), DENV3 (ACY70771.1), and DENV4 (AEW50183.1). |  |  |  |

**Figure 5.**
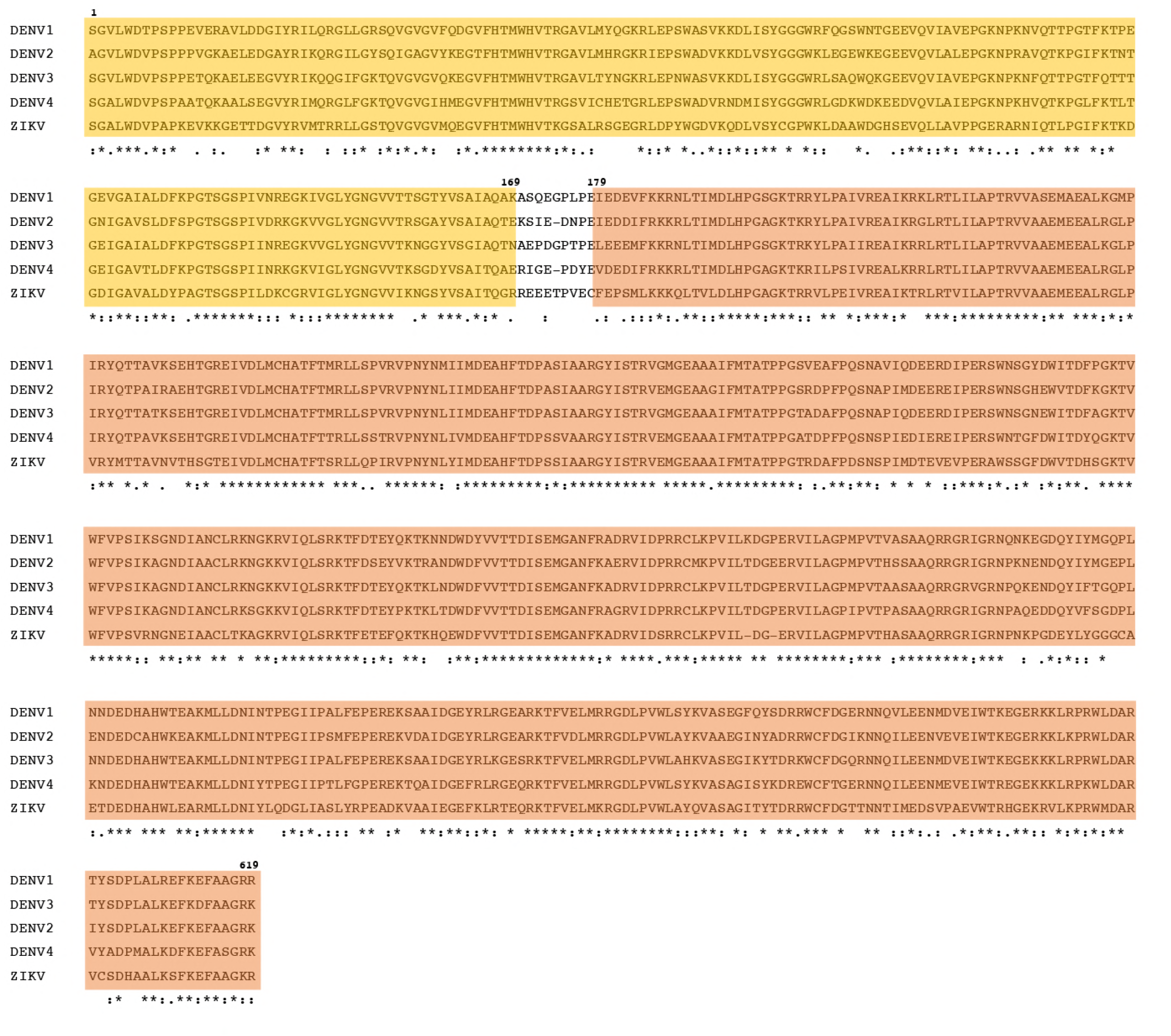
Clustal Omega generated amino acid sequence alignment of DENV serotypes 1 to 4 and Asian ZIKV. The residues in yellow represent the NS3 protease domain (amino acids 1-169) and the residues in orange represent the helicase domain (amino acids 179-619). *, single, fully conserved residue. :, conservation between groups of strongly similar properties – scoring > 0.5 in the Gonnet PAM 250 matrix., conservation between groups of weakly similar properties – scoring =< 0.5 in the Gonnet PAM 250 matrix.

A relatively large body of epidemiological and laboratory-based evidence has suggested that severe and often fatal forms of dengue disease occurs most commonly during a secondary infection by a heterotypic DENV serotype (29, 30). Another phenomenon, known as original antigenic sin (OAS), has been observed in antibody as well as T-cell responses, in which less-effective T cells generated in response to a primary DENV infection predominate during a subsequent infection with a different DENV serotype, resulting in an inappropriate response and predisposing individuals to severe disease (31, 32). The OAS hypothesis was challenged by a study in Sri Lankan individuals infected with DENV, which showed that the phenomenon does not generate less functional responses, but instead correlates with protective responses to conserved viral sequences (26). Unexpectedly, we did not observe T cell responses that were significantly higher in magnitude in individuals with prior DENV exposure. These results are in contrast to our data on African ZIKV infections, which showed that previous flavivirus exposure was associated with enhanced T cell responses (27). One possibility is that the proportion of HIV infection among those with prior DENV exposure was higher compared with DENV-naïve in this study (90.5% versus 50% comparing sDENV and pDENV cases; 83.3% versus 60% comparing ZIKVwpDENV and pZIKV cases). Nevertheless, as co-circulation of DENV, ZIKV, and other flaviviruses occurs throughout many parts of the world, it is critical to continue to develop tools to better understand T cell immunity in individuals exposed to multiple flaviviruses.

A recent study using human leukocyte antigen (HLA) transgenic mice infected with DENV2 and Asian and African ZIKV strains revealed cross-reactive T cell responses to HLA-restricted epitopes (25). Out of 8 ZIKV NS3 epitopes computationally predicted to bind HLA class I molecules, only 3 epitopes elicited DENV2/Asian ZIKV cross-reactive T cell responses. Of note, the cross-reactive epitopes were all positioned within the helicase domain of NS3, further supporting our observations of high DENV/ZIKV NS3 helicase T cell cross-reaction. Another study demonstrated ZIKV-specific and ZIKV/DENV cross-reactive T cell responses in humans (10). T cell responses generated in response to prior DENV exposure recognized peptides sequences located throughout the ZIKV proteome. DENV serostatus also influenced T cell immunity to ZIKV. DENV-naïve ZIKV-positive individuals had predominant CD8 T cell responses directed against structural proteins. In contrast, a majority of CD8 T cells responses were directed against nonstructural proteins in DENV-immune ZIKV-positive individuals, suggesting that previous DENV exposure can alter the T cell response.

While the above studies used peptide stimulation to characterize the T cell response, there are concerns around this approach (33). Some of the HLA-predicted peptides may fail to stimulate strong T cell responses as expected. Longer and shorter peptides have also been shown to elicit different types of responses (34–36). An alternative to peptide stimulation is the anthrax LFn, which has the capability to deliver full length antigen into the cytosol for native processing via the MHC pathways, and to elicit better T cell responses compared to peptides in some cases (37–42). Our adaptation of the LFn ELISPOT, not only allowed detection of human DENV and ZIKV infections, but also characterization of the associated T cell responses to structural and nonstructural proteins. We demonstrated that individuals with pDENV and sDENV infections had similar IFN-γ and TNF-α T cell response patterns with high crossreactivity to ZIKV NS3 helicase, but low cross-reactivity to the ZIKV structural proteins. A small number of individuals with sDENV infections had cross-reactive T cell responses to ZIKV structural proteins. Interestingly, however, individuals with pZIKV and ZIKVwpDENV infections had similar IFN-γ and TNF-α T cell responses patterns, with strong responses to structural and nonstructural proteins. It is noteworthy that we observed comparably strong T cell responses to the structural proteins in pZIKV and ZIKVwpDENV cases, in contrast to Grifoni et al., suggesting that the most recent infection may dictate the T cell response (10). Another possibility that cannot be excluded is the differences in T cell stimulation strategies, which may be contributing to the observed differences. Additionally, due to the limited collection of blood samples from each patient, we were unable to characterize CD4- and CD8-specific responses. Future characterization studies using the LFn delivery system on CD4 and CD8 T cells will be important.

Our study of ZIKV seroprevalence in West Africa demonstrated continued human transmission of the virus in HIV- and malaria-infected individuals (43). The co-infection of flaviviruses with HIV or malaria could potentially impact pathophysiological mechanisms, induce different clinical and laboratory findings, and interfere with treatment. Previous studies have shown a suppression of HIV-1 replication during acute DENV infection (44, 45). In this study, we demonstrated that DENV-exposed individuals who are HIV-infected had T cell responses that were significantly lower in magnitude compared to HIV-negative individuals except in individuals with pZIKV infections. We also observed that DENV-exposed HIV-infected individuals have T cell responses that were lower in magnitude to ZIKV proteins compared to DENV-exposed HIV-negative individuals. Whether HIV infection in DENV-exposed individuals reduces the ability to induce cross-reactive T cell responses has important implications. More studies with larger sample sizes are needed to increase our limited understanding of the epidemiological and immunopathogenesis interactions of flavivirus exposure in individuals with HIV and other comorbidities.

In summary, despite high sequence homology between DENV and ZIKV, diagnostic assays based on antibodies to NS1 and T cell responses to NS3 protease are effective at distinguishing human infections by these viruses. The LFn ELISPOT assay has enabled direct comparison of T cell characterization in DENV, Asian and Africa ZIKV human infections. As vaccines against DENV and ZIKV are currently being developed, the information generated from these characterization studies are of high relevance. The results of these characterization studies may contribute to the design and development of DENV and ZIKV vaccines and T cell based diagnostics.

## MATERIALS AND METHODS

### Clinical samples and ethical statement

Fifty late convalescent-phase blood samples were obtained from patients at Professor Edgard Santos University Hospital, Federal University of Bahia, Salvador, Brazil. These individuals were suspected ZIKV-infected during the 2015-2016 ZIKV epidemic and their acute-phase sera were screened for ZIKV and DENV antibodies. Late convalescent-phase peripheral blood mononuclear cells (PBMCs) were separated from whole blood in EDTA tubes by Ficoll-Hypaque gradient density (Sigma-Aldrich, St. Louis, MO, USA) and cryopreserved in freezing media (10% dimethyl sulfoxide [DMSO], Sigma-Aldrich, St. Louis, MO, USA) at −80°C overnight prior to transfer to liquid nitrogen. Convalescent-phase serum was aliquoted and immediately transferred to −80°C.

The Federal University of Bahia Institutional Review Board (IRB), the Harvard T.H. Chan School of Public Health IRB, and the University of Hawaii IRB approved the primary studies under which the samples and data were collected. All patients provided informed consent for the collection of samples. Excess samples and corresponding data were banked, coded prior to analyses, and stored at the Federal University of Bahia.

### ELISAs

For acute-phase sera, commercial ZIKV-NS1 and DENV-E based IgG ELISAs (Euroimmun, Luebeck, Germany) were performed (27). For late convalescent-phase sera, ZIKV- and DENV1-NS1 IgG ELISAs were performed as described previously (22). Briefly, purified NS1 proteins (16 ng per well) were coated onto 96-well plates overnight, followed by blocking and incubation with primary (serum at 1:400 dilution) and secondary (antihuman IgG conjugated with HRP, Jackson) antibodies (22). The OD at 450 nm was read with a reference wavelength of 650 nm. Each ELISA plate included two positives (two confirmed-Zika or confirmed-dengue samples for ZIKV- and DENV-NS1 ELISAs, respectively), four negatives (4 flavivirus-naïve sera), and tested samples (all in duplicates). The OD values were divided by the mean OD value of positive controls to calculate the rOD values. The cut-off was defined by the mean rOD value of negatives plus 12 standard deviations as described previously (22). For samples positive for both ZIKV- and DENV-NS1 ELISAs, the ratio of rOD (=rOD of ZIKV-NS1/rOD of DENV-NS1) was calculated; rOD ratio < or ≥ 0.24 indicated sDENV or ZIKVwpDENV infection, respectively (22).

E protein-based IgG ELISAs using DENV1 virion or ZIKV (MR766 strain) virus like particles (VLP) were also tested for late convalescent-phase sera (46). Briefly, DENV1 virions or ZIKV-VLP derived from ultracentrifugation of culture supernatants of virus-infected Vero cells or pENTR-ZIKV prME plasmid-transfected 293T cells, respectively, were UV inactivated (for virions) and coated on 96-well plates at 4°C overnight, followed by blocking and incubation with primary (serum at 1:800 dilution) and secondary antibodies as above. The rOD and cut-off rOD values were similarly calculated. The difference in rOD of ZIKV and DENV E proteins (ΔrOD=rOD of ZIKV – rOD of DENV) was determined; ΔrOD ≥ 0.17 or < −0.17 was classified as pZIKV or pDENV infection, respectively.

### Neutralization test

PRNT was performed on acute-phase sera to detect neutralization antibody to ZIKV as reported previously (47). For late convalescent-phase sera, a previously described micro-neutralization test was performed (48). Briefly, flat-bottom 96-well plates were seeded with Vero cells (3 × 10^4^ cells per well) 24 h prior to infection. Fourfold serial dilutions of serum (starting from 1:10) were mixed with 50 focus-forming units of DENV1 (Hawaii strain), DENV2 (NGC strain), DENV3 (CH53489), DENV4 (H241 strain) or ZIKV (PRVABC59 strain) at 37°C for 1 h. The mixtures were added to each well followed by incubation for 48 h (except 70 h for DENV1), removal of medium, and fixation as described previously (46). After adding murine mAb 4G2 and secondary antibody mixture (IRDye^®^ 800CW-conjugated goat anti-mouse IgG at 1:10000 and DRAQ5™ Fluorescent Probe at 1:10000), the signal (800 nm/700 nm fluorescence) was detected by Li Cor Odyssey classic (LiCor Biosciences) and analyzed by Image Studio software to determine percent neutralization at different concentrations and NT_90_ as described previously (46, 48).

### LFn fusion protein design

Commercially synthesized gene fragments encoding the NS3 protease and helicase of DENV2 and C, prM, and NS3 protease and helicase of Asian ZIKV and were cloned into the LFn expression vector (pET15bLFn). The pET15bLFn vector contains a T7 promoter, histidine tag (His_6_), and the terminal domain of the anthrax lethal factor (LFn; 255 amino acids). The pET15bLFn containing the coding sequences of the DENV and Asian ZIKV proteins were transformed into *E. coli* BLR (DE3) (Millipore, Medford, MA, USA). Selected clones were sequences to verify the reading frame, and clones containing the correct sequence were used for protein expression.

The LFn-DENV and -ZIKV fusion proteins and the LFn control were expressed upon isopropylthiogalactoside ([IPTG], Sigma-Aldrich, St. Louis, MO, USA) induction in 5L Luria broth containing carbenicillin and chloramphenicol for 2-4 hours. Cells were pelleted by centrifugation and resuspended in imidazole (1mM) binding buffer (Novagen, Madison, WI, USA) in the presence of a protease inhibitor cocktail (Thermo Fisher Scientific, Rockford, IL, USA). Cell pellets were sonicated, centrifuged at 4°C, and the supernatants were loaded in an equilibrated nickel-charged column for affinity purification. The bound proteins were eluted in 100-200 mM imidazole, desalted with a Sephadex G-25M column (Sigma-Aldrich, St. Louis, MO, USA), and eluted in PBS (Sigma-Aldrich, St. Louis, MO, USA). The PBS-eluted proteins were passed through Detoxi-Gel (Thermo Fisher Scientific, Rockford, IL, USA). Protein concentrations were determined samples were stored at −80°C.

### ELISPOT assay

*Ex vivo* ELISPOTs were performed as previously described. Briefly, 96-well polyvinylidene difluoride (PVDF)-backed MultiScreen_HTS_ (MSIP) microtiter plates (Millipore, Medford, MA, USA) were treated with 100ul of 90% ethanol for 30 seconds and washed 5 times with sterile PBS. Plates were coated with 100ul of capture antibodies (Abs) in PBS. Plates containing capture Abs were incubated overnight at 4°C. Plates were then blocked with 1% bovine serum albumin ([BSA], Sigma-Aldrich, St. Louis, MO, USA) in PBS and washed 6 times with PBS. Cryopreserved PBMCs were thawed in R10 medium and incubated overnight at 37°C. PBMCs were washed 2 times with PBS and seeded at 2 × 10^5^ cells/well in a final volume of 100ul/well. LFn-DENV and -ZIKV proteins were added to each well. As a positive control, PBMCs were stimulated with phytohemagglutinin ([PHA], Sigma-Aldrich, St. Louis, MO, USA). As a negative control, wells received LFn. After incubation for 24-28 hours at 37°C in 5% CO_2_, the cells were discarded and plates were washed 3 times with PBS and 3 times with PBS with 0.05% Tween-20 ([PBST], Bio Rad Technologies, Hercules, CA, USA) to remove cells. The detection antibodies were added and plates were incubated overnight at 4°C. Plates were then washed 6 times with PBST, then incubated for 2 hours at room temperature with mixtures containing the enzymatic conjugates. To develop spots, plates were washed 4 times with PBST, three times with PBS, and 1 time with water. Vector Blue substrate solution (Vector Laboratories, Burlingame, CA, USA) was added for 5-15 mins before rinsing with water and air-drying. Digitized images were analyzed for spots using CTL ImmunoSpot reader (Cellular Technology Limited, Cleveland, OH, USA). DENV and ZIKV spots were calculated by subtracting the mean of the negative control value from the mean value of the specific stimulation. Positive responses had to be greater than 4 times the mean background, 3 standard deviations above the background, and ≥55 spot-forming cells per (SFC)/10^6^ PBMCs.

### ROC analysis

The ELISPOTs were validated using PBMCs from individuals that were confirmed DENV- and/or ZIIV-infected by ELISA and/or neutralization tests. DENV and ZIKV NS3 protease to helicase values were calculated, resulting in normalized test ratios (ZIKV NS3 protease divided by DENV NS3 protease) ranging from 0.15-2.95. On the basis of these data, we determined the optimal cutoffs between 0.15 and 2.95 by calculating the sensitivity (number of true positives divided by total confirmed positive values) and specificity (number of true negatives divided by the total confirmed negatives) at increasing 0.05 to the theoretical cutoffs. After calculating the sensitivity and specificity values, the optimal cutoffs were defined as the highest sum of sensitivity and specificity, such that the optimal cutoff values reflected the optimal sensitivity and specificity. The optimal cutoffs obtained for the IFN-γ and TNF-α was 1.05 and 1.048, respectively (Prism 7, GraphPad Software, San Diego, CA, USA).

### Multiple sequence alignment and percent homology analysis

Multiple sequence alignment of DENV1-4 and ZIKV NS3 was performed using the Clustal Omega program (EMBL-EB, Cambridgeshire, UK). Averages of DENV and ZIKV NS3 protease and helicase proteins were calculated using the ExPASy Bioinformatics Resource Portal (Swiss Institute of Bioinformatics, Lausanne, Switzerland) and based on averages of the different homology values in the four DENV serotypes and ZIKV. Average conservation was determined on a per-residue basis for NS3 protease, helicase, and full-length protein.

### Statistical analysis

Statistical analysis was performed using Prism 7 (GraphPad Software, San Diego, CA, USA). Where appropriate, data were expressed as geometric positive means on box whisker and bar graphs ± standard deviation. Data comparisons were conducted using the Wilcoxon rank sum test. A threshold of p<0.05 was considered statistically significant.

### Data Availability

All relevant data has been included in the manuscript. We will provide any additional data upon request.

## ACKNOWLEDGEMENTS

We thank Yichen Lu for providing us with the LFn expression vector and Gwong-Jen J. Chang at the CDC Fort Collins for providing us the pENTR-ZIKV prME plasmid. This work was funded by a Harvard University David Rockefeller Center for Latin American Studies grant to BBH, and by grants R01AI110769-01 from the National Institute of Allergy and Infectious Diseases and P20GM103516 from the National Institute of General Medical Sciences, NIH to WKW. The funders had no role in study design, data collection and analysis, decision to publish, or preparation of the manuscript.

## REFERENCES

1. Peirson TC, Diamond MS. 2013. Flaviviruses. Fields Virology, 6th ed. Lippincott Williams & Wilkins, Philadelphia.

2. Bhatt S, Gething PW, Brady OJ, Messina JP, Farlow AW, Moyes CL, Drake JM, Brownstein JS, Hoen AG, Sankoh O, Myers MF, George DB, Jaenisch T, Wint GR, Simmons CP, Scott TW, Farrar JJ, Hay SI. 2013. The global distribution and burden of dengue. Nature 496:504–7.

3. Faye O, Freire CC, Iamarino A, Faye O, de Oliveira JV, Diallo M, Zanotto PM, Sall AA. 2014. Molecular evolution of Zika virus during its emergence in the 20(th) century. PLoS Negl Trop Dis 8:e2636.

4. Simpson DI. 1964. Zika Virus Infection in Man. Trans R Soc Trop Med Hyg 58:335–8.

5. Weaver SC, Costa F, Garcia-Blanco MA, Ko AI, Ribeiro GS, Saade G, Shi PY, Vasilakis N. 2016. Zika virus: History, emergence, biology, and prospects for control. Antiviral Res 130:69–80.

6. Krauer F, Riesen M, Reveiz L, Oladapo OT, Martinez-Vega R, Porgo TV, Haefliger A, Broutet NJ, Low N, Group WHOZCW. 2017. Zika Virus Infection as a Cause of Congenital Brain Abnormalities and Guillain-Barre Syndrome: Systematic Review. PLoS Med 14:e1002203.

7. WorldHealth Organization. 2016. WHO Director-General summarizes the outcome of the Emergency Committee regarding clusters of microcephaly and Guillain-Barré syndrome. http://www.who.int/mediacentre/news/statements/2016/emergency-committee-zika-microcephaly/en/. Accessed

8. Armstrong P, Hennessey M, Adams M, Cherry C, Chiu S, Harrist A, Kwit N, Lewis L, McGuire DO, Oduyebo T, Russell K, Talley P, Tanner M, Williams C, Zika Virus Response E, Laboratory T. 2016. Travel-Associated Zika Virus Disease Cases Among U.S. Residents–United States, January 2015-February 2016. MMWR Morb Mortal Wkly Rep 65:286–9.

9. Waggoner JJ, Gresh L, Vargas MJ, Ballesteros G, Tellez Y, Soda KJ, Sahoo MK, Nunez A, Balmaseda A, Harris E, Pinsky BA. 2016. Viremia and Clinical Presentation in Nicaraguan Patients Infected With Zika Virus, Chikungunya Virus, and Dengue Virus. Clin Infect Dis 63:1584–1590.

10. Grifoni A, Pham J, Sidney J, O’Rourke PH, Paul S, Peters B, Martini SR, de Silva AD, Ricciardi MJ, Magnani DM, Silveira CGT, Maestri A, Costa PR, de-Oliveira-Pinto LM, de Azeredo EL, Damasco PV, Phillips E, Mallal S, de Silva AM, Collins M, Durbin A, Diehl SA, Cerpas C, Balmaseda A, Kuan G, Coloma J, Harris E, Crowe JE, Jr., Stone M, Norris PJ, Busch M, Vivanco-Cid H, Cox J, Graham BS, Ledgerwood JE, Turtle L, Solomon T, Kallas EG, Watkins DI, Weiskopf D, Sette A. 2017. Prior Dengue virus exposure shapes T cell immunity to Zika virus in humans. J Virol doi:10.1128/JVI.01469-17.

11. Lanciotti RS, Kosoy OL, Laven JJ, Velez JO, Lambert AJ, Johnson AJ, Stanfield SM, Duffy MR. 2008. Genetic and serologic properties of Zika virus associated with an epidemic, Yap State, Micronesia, 2007. Emerg Infect Dis 14:1232–9.

12. Tsai WY, Youn HH, Brites C, Tsai JJ, Tyson J, Pedroso C, Drexler JF, Stone M, Simmons G, Busch MP, Lanteri M, Stramer SL, Balmaseda A, Harris E, Wang WK. 2017. Distinguishing Secondary Dengue Virus Infection From Zika Virus Infection With Previous Dengue by a Combination of 3 Simple Serological Tests. Clin Infect Dis 65:1829–1836.

13. Lai CY, Tsai WY, Lin SR, Kao CL, Hu HP, King CC, Wu HC, Chang GJ, Wang WK. 2008. Antibodies to envelope glycoprotein of dengue virus during the natural course of infection are predominantly cross-reactive and recognize epitopes containing highly conserved residues at the fusion loop of domain II. J Virol 82:6631–43.

14. Johnson BW, Kosoy O, Martin DA, Noga AJ, Russell BJ, Johnson AA, Petersen LR. 2005. West Nile virus infection and serologic response among persons previously vaccinated against yellow fever and Japanese encephalitis viruses. Vector Borne Zoonotic Dis 5:137–45.

15. Rabe IB, Staples JE, Villanueva J, Hummel KB, Johnson JA, Rose L, Mts, Hills S, Wasley A, Fischer M, Powers AM. 2016. Interim Guidance for Interpretation of Zika Virus Antibody Test Results. MMWR Morb Mortal Wkly Rep 65:543–6.

16. Centers for Disease Control and Prevention. Guidance for U.S. laboratories testing for Zika virus infection. Available at: http://www.cdc.gov/zika/laboratories/lab-guidance.html. Accessed 14 February 2018.

17. Dejnirattisai W, Supasa P, Wongwiwat W, Rouvinski A, Barba-Spaeth G, Duangchinda T, Sakuntabhai A, Cao-Lormeau VM, Malasit P, Rey FA, Mongkolsapaya J, Screaton GR. 2016. Dengue virus sero-cross-reactivity drives antibody-dependent enhancement of infection with zika virus. Nat Immunol 17:1102–8.

18. Priyamvada L, Quicke KM, Hudson WH, Onlamoon N, Sewatanon J, Edupuganti S, Pattanapanyasat K, Chokephaibulkit K, Mulligan MJ, Wilson PC, Ahmed R, Suthar MS, Wrammert J. 2016. Human antibody responses after dengue virus infection are highly cross-reactive to Zika virus. Proc Natl Acad Sci U S A 113:7852–7.

19. Castanha PM, Braga C, Cordeiro MT, Souza AI, Silva CD, Jr., Martelli CM, van Panhuis WG, Nascimento EJ, Marques ET. 2016. Placental Transfer of Dengue Virus (DENV)-Specific Antibodies and Kinetics of DENV Infection-Enhancing Activity in Brazilian Infants. J Infect Dis 214:265–72.

20. Stettler K, Beltramello M, Espinosa DA, Graham V, Cassotta A, Bianchi S, Vanzetta F, Minola A, Jaconi S, Mele F, Foglierini M, Pedotti M, Simonelli L, Dowall S, Atkinson B, Percivalle E, Simmons CP, Varani L, Blum J, Baldanti F, Cameroni E, Hewson R, Harris E, Lanzavecchia A, Sallusto F, Corti D. 2016. Specificity, cross-reactivity, and function of antibodies elicited by Zika virus infection. Science 353:823–6.

21. Halstead SB, O’Rourke EJ. 1977. Antibody-enhanced dengue virus infection in primate leukocytes. Nature 265:739–41.

22. Bardina SV, Bunduc P, Tripathi S, Duehr J, Frere JJ, Brown JA, Nachbagauer R, Foster GA, Krysztof D, Tortorella D, Stramer SL, Garcia-Sastre A, Krammer F, Lim JK. 2017. Enhancement of Zika virus pathogenesis by preexisting antiflavivirus immunity. Science 356:175–180.

23. Steinhagen K, Probst C, Radzimski C, Schmidt-Chanasit J, Emmerich P, van Esbroeck M, Schinkel J, Grobusch MP, Goorhuis A, Warnecke JM, Lattwein E, Komorowski L, Deerberg A, Saschenbrecker S, Stocker W, Schlumberger W. 2016. Serodiagnosis of Zika virus (ZIKV) infections by a novel NS1-based ELISA devoid of cross-reactivity with dengue virus antibodies: a multicohort study of assay performance, 2015 to 2016. Euro Surveill 21.

24. Balmaseda A, Stettler K, Medialdea-Carrera R, Collado D, Jin X, Zambrana JV, Jaconi S, Cameroni E, Saborio S, Rovida F, Percivalle E, Ijaz S, Dicks S, Ushiro- Lumb I, Barzon L, Siqueira P, Brown DWG, Baldanti F, Tedder R, Zambon M, de Filippis AMB, Harris E, Corti D. 2017. Antibody-based assay discriminates Zika virus infection from other flaviviruses. Proc Natl Acad Sci U S A 114:8384–8389.

25. Wen J, Tang WW, Sheets N, Ellison J, Sette A, Kim K, Shresta S. 2017. Identification of Zika virus epitopes reveals immunodominant and protective roles for dengue virus cross-reactive CD8(+) T cells. Nat Microbiol 2:17036.

26. Weiskopf D, Angelo MA, de Azeredo EL, Sidney J, Greenbaum JA, Fernando AN, Broadwater A, Kolla RV, De Silva AD, de Silva AM, Mattia KA, Doranz BJ, Grey HM, Shresta S, Peters B, Sette A. 2013. Comprehensive analysis of dengue virus-specific responses supports an HLA-linked protective role for CD8+ T cells. Proc Natl Acad Sci U S A 110:E2046–53.

27. Herrera BB, Tsai WY, Chang CA, Hamel DJ, Wang WK, Lu Y, Mboup S, Kanki PJ. 2018. Sustained specific and cross-reactive T cell responses to Zika and Dengue viruses NS3 in West Africa. J Virol doi:10.1128/JVI.01992-17.

28. Netto EM, Moreira-Soto A, Pedroso C, Hoser C, Funk S, Kucharski AJ, Rockstroh A, Kummerer BM, Sampaio GS, Luz E, Vaz SN, Dias JP, Bastos FA, Cabral R, Kistemann T, Ulbert S, de Lamballerie X, Jaenisch T, Brady OJ, Drosten C, Sarno M, Brites C, Drexler JF. 2017. High Zika Virus Seroprevalence in Salvador, Northeastern Brazil Limits the Potential for Further Outbreaks. MBio 8.

29. Halstead SB. 2007. Dengue. Lancet 370:1644–52.

30. Guzman MG, Alvarez M, Halstead SB. 2013. Secondary infection as a risk factor for dengue hemorrhagic fever/dengue shock syndrome: an historical perspective and role of antibody-dependent enhancement of infection. Arch Virol 158:144559.

31. Halstead SB, Rojanasuphot S, Sangkawibha N. 1983. Original antigenic sin in dengue. Am J Trop Med Hyg 32:154–6.

32. Mongkolsapaya J, Dejnirattisai W, Xu XN, Vasanawathana S, Tangthawornchaikul N, Chairunsri A, Sawasdivorn S, Duangchinda T, Dong T, Rowland-Jones S, Yenchitsomanus PT, McMichael A, Malasit P, Screaton G. 2003. Original antigenic sin and apoptosis in the pathogenesis of dengue hemorrhagic fever. Nat Med 9:921–7.

33. Karlsson RK, Jennes W, Page-Shafer K, Nixon DF, Shacklett BL. 2004. Poorly soluble peptides can mimic authentic ELISPOT responses. J Immunol Methods 285:89–92.

34. Madden DR. 1995. The three-dimensional structure of peptide-MHC complexes. Annu Rev Immunol 13:587–622.

35. Stryhn A, Pedersen LO, Holm A, Buus S. 2000. Longer peptide can be accommodated in the MHC class I binding site by a protrusion mechanism. Eur J Immunol 30:3089–99.

36. Horig H, Young AC, Papadopoulos NJ, DiLorenzo TP, Nathenson SG. 1999. Binding of longer peptides to the H-2Kb heterodimer is restricted to peptides extended at their C terminus: refinement of the inherent MHC class I peptide binding criteria. J Immunol 163:4434–41.

37. Cao H, Agrawal D, Kushner N, Touzjian N, Essex M, Lu Y. 2002. Delivery of exogenous protein antigens to major histocompatibility complex class I pathway in cytosol. J Infect Dis 185:244–51.

38. Kushner N, Zhang D, Touzjian N, Essex M, Lieberman J, Lu Y. 2003. A fragment of anthrax lethal factor delivers proteins to the cytosol without requiring protective antigen. Proc Natl Acad Sci U S A 100:6652–7.

39. Lu Y, Friedman R, Kushner N, Doling A, Thomas L, Touzjian N, Starnbach M, Lieberman J. 2000. Genetically modified anthrax lethal toxin safely delivers whole HIV protein antigens into the cytosol to induce T cell immunity. Proc Natl Acad Sci U S A 97:8027–32.

40. McEvers K, Elrefaei M, Norris P, Deeks S, Martin J, Lu Y, Cao H. 2005. Modified anthrax fusion proteins deliver HIV antigens through MHC Class I and II pathways. Vaccine 23:4128–35.

41. Sarr AD, Lu Y, Sankale JL, Eisen G, Popper S, Mboup S, Kanki PJ, Cao H. 2001. Robust HIV type 2 cellular immune response measured by a modified anthrax toxin-based enzyme-linked immunospot assay. AIDS Res Hum Retroviruses 17:1257–64.

42. Liao Q, Strong AJ, Liu Y, Liu Y, Meng P, Fu Y, Touzjian N, Shao Y, Zhao Z, Lu Y. 2012. HIV vaccine candidates generate in vitro T cell response to putative epitopes in Chinese-origin rhesus macaques. Vaccine 30:1601–8.

43. Herrera BB, Chang CA, Hamel DJ, Mboup S, Ndiaye D, Imade G, Okpokwu J, Agbaji O, Bei AK, Kanki PJ. 2017. Continued Transmission of Zika Virus in Humans in West Africa, 1992-2016. J Infect Dis 215:1546–1550.

44. Watt G, Kantipong P, Jongsakul K. 2003. Decrease in human immunodeficiency virus type 1 load during acute dengue fever. Clin Infect Dis 36:1067–9.

45. McLinden JH, Stapleton JT, Chang Q, Xiang J. 2008. Expression of the dengue virus type 2 NS5 protein in a CD4(+) T cell line inhibits HIV replication. J Infect Dis 198:860–3.

46. Tsai WY, Lai CY, Wu YC, Lin HE, Edwards C, Jumnainsong A, Kliks S, Halstead S, Mongkolsapaya J, Screaton GR, Wang WK. 2013. High-avidity and potently neutralizing cross-reactive human monoclonal antibodies derived from secondary dengue virus infection. J Virol 87:12562–75.

47. Moreira-Soto A, Sarno M, Pedroso C, Netto EM, Rockstroh A, Luz E, Feldmann M, Fischer C, Bastos FA, Kummerer BM, de Lamballerie X, Drosten C, Ulbert S, Brites C, Drexler JF. 2017. Evidence for Congenital Zika Virus Infection From Neutralizing Antibody Titers in Maternal Sera, Northeastern Brazil. J Infect Dis 216:1501–1504.

48. Govindarajan D, Meschino S, Guan L, Clements DE, ter Meulen JH, Casimiro DR, Coller BA, Bett AJ. 2015. Preclinical development of a dengue tetravalent recombinant subunit vaccine: Immunogenicity and protective efficacy in nonhuman primates. Vaccine 33:4105–16.

